# Integrated Likelihood for Phylogenomics under a No-Common-Mechanism Model

**DOI:** 10.1101/500520

**Authors:** H. Tidwell, L. Nakhleh

## Abstract

The availability of genome-wide sequence data from a large number of species as well as data from multiple individuals within a species has ushered in the era of phylogenomics. In this era, species phylogeny inference is based on models of sequence evolution on gene trees as well as models of gene tree evolution within the branches of species phylogenies. Parsimony, likelihood, Bayesian, and distance methods have been introduced for species phylogeny inference based on such models. All methods, except for the parsimony ones, assume a common mechanism across all loci as captured by a single value of each branch length of the species phylogeny. In this paper, we propose a “no common mechanism” (NCM) model, where every gene tree evolves according to its own parameters of the species phylogeny. An analogous model was proposed and explored, both mathematically and experimentally, for sites, or characters, in a sequence alignment in the context of the classical phylogeny problem. For example, a famous equivalence between the maximum parsimony and maximum likelihood phylogeny estimates was established under certain NCM models by Tuffley and Steel. Here we derive an analytically integrated likelihood of both species trees and networks given the gene trees of multiple loci under an NCM model. We demonstrate the performance of inference under this integrated likelihood on both simulated and biological data. The model presented here will afford opportunities for exploring connections among various methods for estimating species phylogenies from multiple, independent loci.

## 1 Introduction

A phylogenetic tree models the evolutionary history of a set of taxa (genes, species, etc.) from their most recent common ancestor. Analyses of genome-wide data from several groups of species have highlighted a significant phenomenon, namely the incongruence among phylogenetic trees of the different genomic regions as well as with the phylogeny of the species [8]. For example, Pollard *et al*. showed that all three possible rooted gene tree topologies are supported by different regions across the genomes of three *Drosophila* species, only one of which, of course, agrees with the topology of the species tree [11]. Similarly, by analyzing the genomes of humans, chimpanzees, and gorillas, Scally *et al*. showed that while the species tree has human and chimpanzee sharing a most recent common ancestor before either of them shares a common ancestor with gorilla, 30% of the genome places gorilla closer to human or chimpanzee than the latter are to each other [13]. In both cases, the authors named *incomplete lineage sorting* (ILS) as the main cause of incongruence among the gene trees of the different genomic regions. ILS is mathematically well understood under the *multispecies coalescent* (MSC) model [2]. Fig. 1 illustrates the gene tree probability distribution that the multispecies coalescent defines.

**Figure 1:**
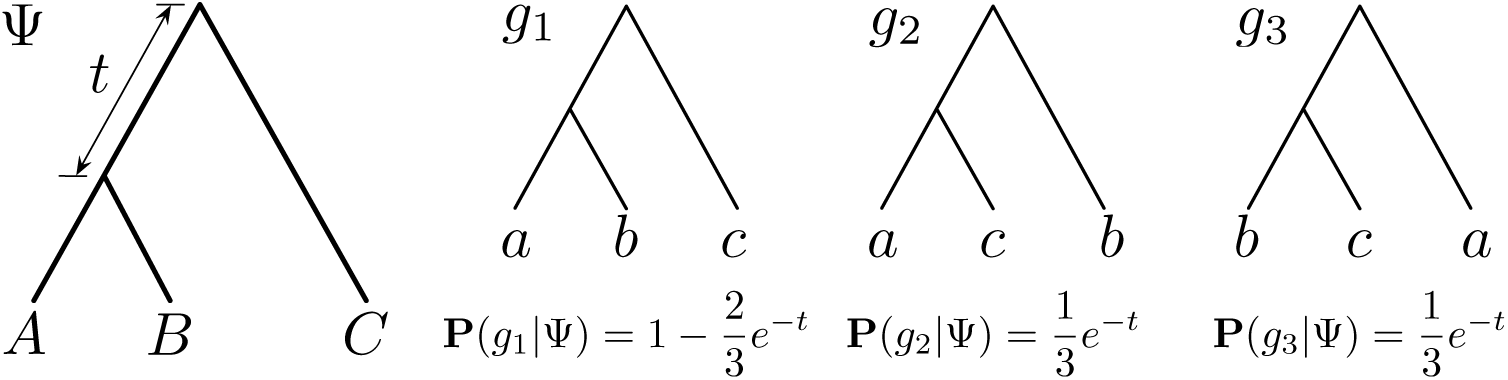
The multispecies coalescent (MSC) model. The species tree Ψ defines a probability distribution on gene tree topologies. Assuming individuals *a*, *b*, and *c* are sampled from species *A*, *B*, and *C*, respectively, the three distinct rooted gene tree topologies, *g*_1_, *g*_2_, and *g*_3_, are shown, along with their probabilities under MSC. The branch length *t* here is given in coalescent units (one coalescent unit in this case equals 2*N* generations, where *N* is the effective population size of the ancestor of *A* and *B*).

However, as Maddison noted [8], other processes could give rise to incongruence among gene trees, including gene duplication and loss, and horizontal gene transfer. In the latter case, the evolutionary history of the genomes is best modeled by a phylogenetic network [9]. The multispecies coalescent has been extended to incorporate such processes [12, 24, 25, 20, 21]. These findings have given rise to phylogenomics—the inference of a species phylogeny from genome-wide data. Given *m* gene trees 𝒢= {*g*_1_,…, *g*_*m*_} for *m* independent loci (genomic regions), the likelihood of a species phylogeny Ψ and its branch lengths *λ* is given by

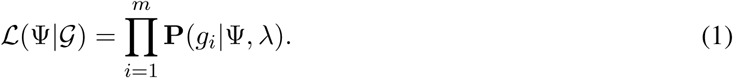

Assuming, for example, that ILS is the sole cause of all incongruence among gene trees in 𝒢, then **P**(*g*_*i*_*|*Ψ, *λ*) is given by MSC, as illustrated in Fig. 1. If both ILS and reticulation are at play, then the probability distribution is given by the multispecies network coalescent [24] and if ILS and duplication/loss are at play, then the probability distribution can be derived based on the model of Rasmussen and Kellis [12].

The formulation given by Eq. (1) assumes a “common mechanism” across all loci—all gene trees “grow” within a species tree given by a single setting of branch lengths. In this paper we explore, for the first time to our knowledge, a “no common mechanism,” or NCM, model, which states that each gene tree evolved within the branches of the species phylogeny under a totally separate process from all other gene trees as given by the parameters (e.g., branch lengths) of the species phylogeny. It is important to note that the NCM model has been explored and studied in the “classical” phylogeny problem (inferring a phylogenetic tree from a sequence alignment). Under that setting, NCM posits that each site in the sequence alignment evolved under its own branch lengths of the phylogenetic tree. Tuffley and Steel [18] established a seminal result in the field by proving that the maximum parsimony and maximum likelihood estimates of a phylogenetic tree are equal under an NCM model based on a symmetric Poisson process of nucleotide substitution. Additional mathematical results based on the NCM model were later established by Steel and Penny [15]. The NCM model allowed for analytically integrating out the branch lengths of a phylogenetic tree and, consequently, efficiently exploring the space of phylogenetic trees [7]. Steel [14] showed that it is possible to achieve statistical consistency of inference under certain NCM models.

However, while the NCM model was used by some as an argument in favor of using maximum parsimony, Holder *et al*. [4] showed problems with this argument. Furthermore, Huelsenbeck *et al*. [6] argued that “biologically inspired phylogenetic models” outperform the NCM model. More specifically, it is hard to justify a phylogenetic model by which every site, including adjacent ones, has its own evolutionary process. This is why the NCM model was deemed more useful for morphological characters than molecular characters in a sequence alignment. The question is: Is the NCM model appropriate in phylogenomics where loci are sampled sufficiently far apart from across the genome? Different loci could be under different selective pressures. Loci on different chromosomes (e.g., autosomes vs. sex chromosomes) could have evolved under different processes. In this paper, we introduce an NCM model for phylogenomics, analytically derive an integrated likelihood model under the multispecies coalescent with the NCM, and show the performance of inference based on this NCM model on data simulated under a common mechanism, as well as a biological data set. As Huelsenbeck *et al*. argued [7], the work we present here could lead to more efficient ways of exploring the space of species phylogenies. Furthermore, this NCM model could give rise to research avenues into connections between maximum likelihood and maximum parsimony inference of species trees from gene trees (in this case, one parsimony formulation is given by the “minimizing deep coalescences” criterion [8, 16, 23]).

## 2 Background

In order to account for both reticulation and incomplete lineage sorting, we use the phylogenetic network model since it generalizes trees.

### Definition 1

*A* phylogenetic *χ*-network, *or χ-network for short*, Ψ *is a rooted*, *directed*, *acyclic graph (DAG) with set of nodes V* (Ψ) = {*r*} ∪ *V*_*L*_ ∪ *V*_*T*_ ∪ *R*, *where*

- *indeg*(*r*) = 0 *(r is the* root *of* Ψ);
- *∀v ∈ V*_*L*_, *indeg*(*v*) = 1 *and outdeg*(*v*) = 0 *(V*_*L*_ *are the* external tree nodes, *or* leaves, *of* Ψ);
- *∀v ∈ V*_*T*_, *indeg*(*v*) = 1 *and outdeg*(*v*) *≥* 2 *(V*_*T*_ *are the* internal tree nodes *of* Ψ*); and*,
- *∀v ∈ R*, *indeg*(*v*) = 2 *and outdeg*(*v*) = 1 *(R are the* reticulation nodes *of* Ψ),

*and set of edges E*(Ψ) ⊆ *V × V that consists of* reticulation edges, *whose heads are reticulation nodes*, *and* tree edges, *whose heads are tree nodes*. *The leaves of the network are bijectively labeled by elements of χ*.

We assume that we have *ℓ* reticulation nodes *R* = {*r*_1_,…, *r*_*ℓ*_} with *ℓ* associated inheritance probabilities *γ*_1_,…, *γ*_*ℓ*_, respectively (that is, node *r*_*i*_ has two parents *pr*_*i*_ and 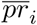 with inheritance probabilities *γ*_*i*_ and (1–*γ*_*i*_) associated with the edges *b*1_*i*_ = (*pr*_*i*_, *r*_*i*_) and 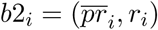, respectively).

In addition to the topology of a phylogenetic network Ψ, each edge *b* = (*u*, *v*) in *E*(Ψ) has a length *λ*_*b*_ measured in coalescent units, which is the number of generations divided by effective population size on that branch. We use Ψ to refer to the topology of the phylogenetic network, Λ to refer to its branch lengths, and Γ to refer to the inheritance probabilities associated with all reticulation nodes. A species tree is a phylogenetic network with no reticulation nodes (and an empty Γ).

### 2.1 Distribution of Gene Tree Topologies

Given a phylogenetic network Ψ, its branch lengths Λ and inheritance probabilities Γ on the reticulation edges, the gene tree topology is a random variable whose probability mass function (pmf) we now briefly review. This pmf was originally derived for the case of species trees by Degnan and Salter [1] and later extended to the case of phylogenetic networks by Yu *et al*. [24].

We denote by Ψ_*u*_ the set of nodes that are reachable from the root of Ψ via at least one path that goes through node *u ∈ V* (Ψ). Then given a phylogenetic network Ψ and a gene tree *g* for some locus *j*, a coalescent history is a function *h*: *V* (*g*) *→ E*(Ψ) such that the following two conditions hold:

- if *v* is a leaf in *g*, then *h*(*v*) = (*x*, *y*) where *y* is the leaf in Ψ with the label of the species from which the allele labeling leaf *v* in *G* is sampled;
- if *v* is a node in the subtree of *g* that is rooted at *u*, and *h*(*u*) = (*p*, *q*), then *h*(*v*) = (*x*, *y*) where *y* ∈ Ψ_*q*_.

Given a phylogenetic network Ψ and a gene tree *g* for locus *j*, we denote by *H*_Ψ_(*g*) the set of all coalescent histories of *g* within the branches of Ψ. Then the pmf of the gene tree is given by

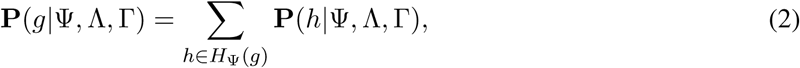

where Λ are the branch lengths of the phylogenetic network (in coalescent units), Γ is the inheritance probabilities matrix, and **P**(*h*| Ψ, Λ, Γ) gives the pmf of the coalescent history random variable, which can be computed as

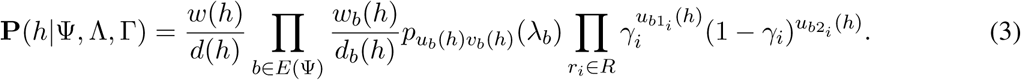

In this equation, *u*_*b*_(*h*) and *v*_*b*_(*h*) denote the number of lineages that respectively enter and exit edge *b* of Ψ under coalescent history *h*. The term 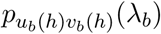 is the probability of *u*_*b*_(*h*) gene lineages coalescing into *v*_*b*_(*h*) during time *λ*_*b*_. And *w*_*b*_(*h*)*/d*_*b*_(*h*) is the proportion of all coalescent scenarios resulting from *u*_*b*_(*h*) – *v*_*b*_(*h*) coalescent events that agree with the topology of the gene tree. This quantity without the *b* subscript corresponds to the root of Ψ. Notice that removing the rightmost product over the reticulation nodes in Eq. (3) gives the pmf for species trees.

## 3 Methods

We now show how to analytically integrate the branch lengths and inheritance probabilities.

### Integrating out the branch lengths

The function *p*_*uv*_(*t*) employed by Eq. (3) is given by

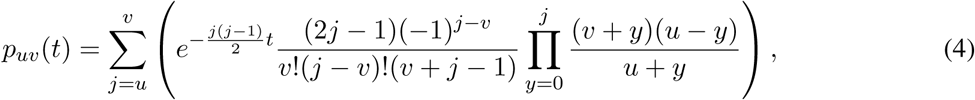

which, for simplifying the equations below, can be written as

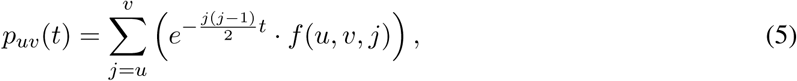

where 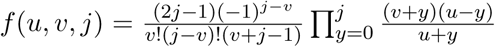 Assuming a truncated Exponential prior with support in (0, *τ*] and hyperparameter value of 1 on *t*, we have

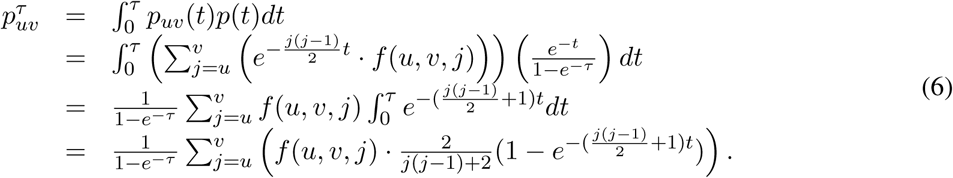

Using this result, we have

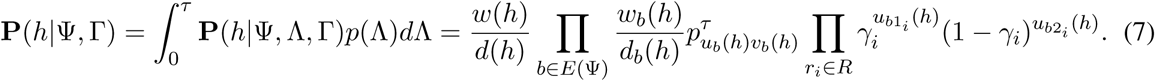

### Integrating out the inheritance probabilities

We now have

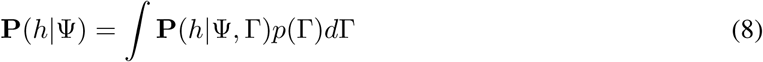

where *p*(Γ) is a prior on the inheritance probabilities, and the multiple integration is taken over all *ℓ* gamma’s on (0, 1). We assume *γ*_*i*_ *∼ Beta*(2, 2), so that we have a conjugate prior (pdf in this case is 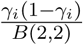. Then, we have

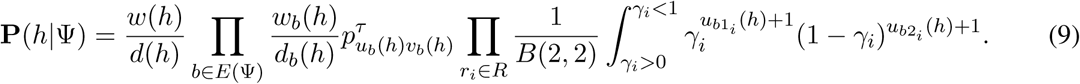

Using the Binomial Theorem, and denoting 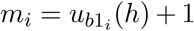 and 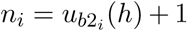 to shorten notation, we have

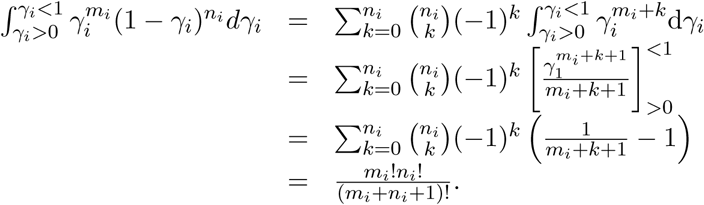

Putting it altogether, we obtain

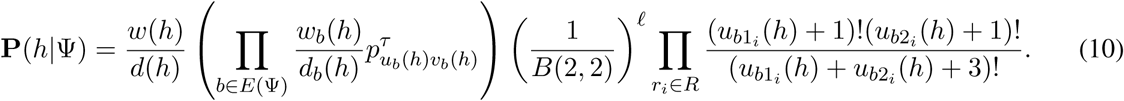

Finally,

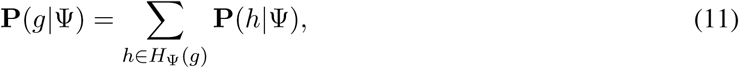

where Ψ is given by its topology alone.

Observe that if one treats the branch lengths in Eq. (1) as a nuisance parameter, then the integrated likelihood is given by

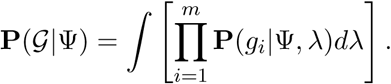

### Inference

The calculation given by Eq. (11) above allows us to compute

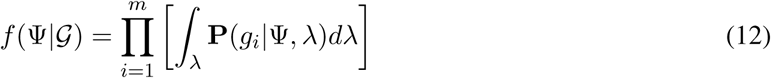

from a set 𝒢 of input gene trees inferred on multiple independent loci. It is unclear, though, under what conditions, if any, both Eq. (1) and Eq. (12) yield the same values. For inferring an optimal network under Eq. (12), also known as the maximum integrated likelihood network, a search for the network Ψ that maximizes *f* (Ψ|𝒢) is conducted. Since no branch lengths or inheritance probabilities are optimized or sampled, just like the case of inference under the MDC criterion, we use the exact search heuristic and moves of [23].

## 4 Results

### 4.1 Evaluating the NCM Criterion on Simulated Data

We first set out to study the performance of maximum integrated likelihood inference under the NCM model, and compare it to that of inference under the parsimony criterion “minimizing deep coalescence” (MDC) of [23]. We follow the same simulation setup, including the model networks, parameters, and numbers of gene trees as that in [23]. More specifically, we considered four phylogenetic network topologies involving distinct combinations of reticulation and speciation events, as shown in Fig. 2. To better understand the effects of deep coalescence in each scenario, we used two settings of the branch length parameters for each network. In branch length setting 1, each of the values *t*1, *t*2, *t*3, and *t*4 are equal to 1 coalescent unit. In branch length setting 2, each value is equal to 2 coalescent units. Setting 1 should involve more deep coalescence events, while setting 2 involves longer branches which are less likely according to the exponential prior on the branch lengths. Each provides unique challenges for the integrated likelihood inference under NCM. For each setting and for each number of loci in the set {10, 25, 50, 100, 500, 1000, 2000}, we generated 100 data sets of gene trees using the program ms [5]. It is important to highlight here that the data was generated under a common mechanism; that is, not under the NCM model. We then ran the inference method under MDC of [23] and the maximum integrated likelihood inference under the NCM model on each data set. We then computed the topological distance of [10], as implemented in PhyloNet [17], between each inferred network and the model network on which the data was generated, and averaged the results over all 100 data sets for each setting. The results are shown in Fig. 3. Note that for the calculation of the NCM integrated likelihood, we used a non-truncated exponential prior for the branch lengths of the species phylogeny. This is equivalent to letting the hyperparameter *τ* grow arbitrarily large, and results in a similar likelihood function as that in (6).

**Figure 2:**
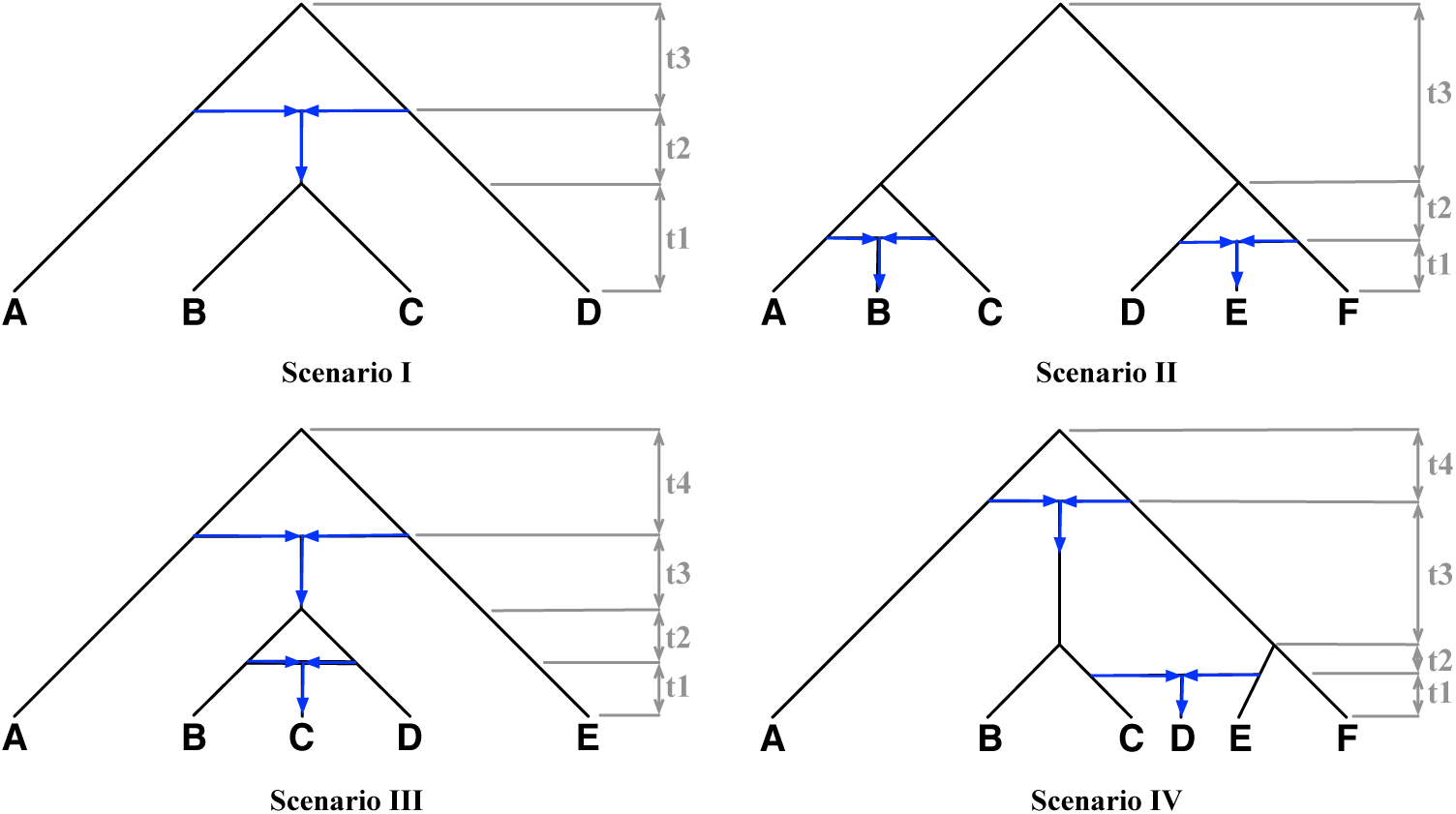
Phylogenetic networks considered in the simulation study. All inheritance probabilities were set to 0.5 in the simulations. Blue arrows indicated directions into and out of the reticulation nodes.

**Figure 3:**
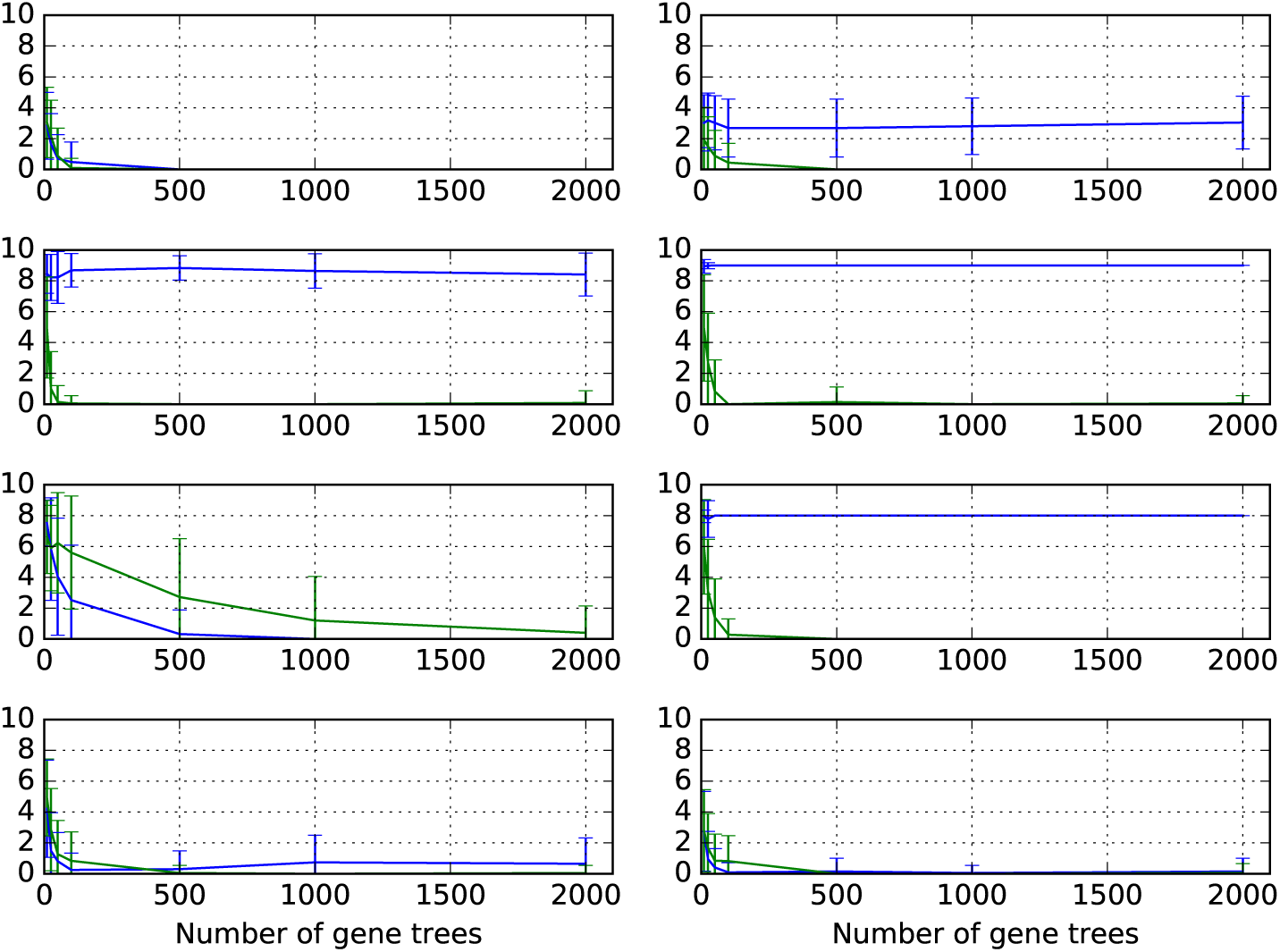
Accuracy of network inference on simulated scenarios. The symmetric network difference between the inferred and model network, averaged over 100 trials, using the MDC criterion as implemented in [23] (green) and the maximum integrated likelihood under the NCM model (blue). Rows from top to bottom correspond to Scenarios I-IV, respectively, of Fig. 2. Left and right columns correspond to branch length settings 1 and 2, respectively.

As the results show, both methods have almost the same behavior and accuracy under branch length setting 1 for the networks of Scenarios I, III, and IV, and under branch length setting 2 for the network of Scenario IV. However, while MDC always converged onto the true network, inference under the NCM model diverged from the true network in the other cases. We highlight again that the data was not generated under the NCM model. Therefore, these results demonstrate the behavior of inference under an NCM model when all loci have an underlying common mechanism.

We then set out to compare the erroneous networks inferred under the NCM model (Fig. 4) to their true counterparts. A quick inspection of the three networks in the figure points to a very interesting pattern.

**Figure 4:**
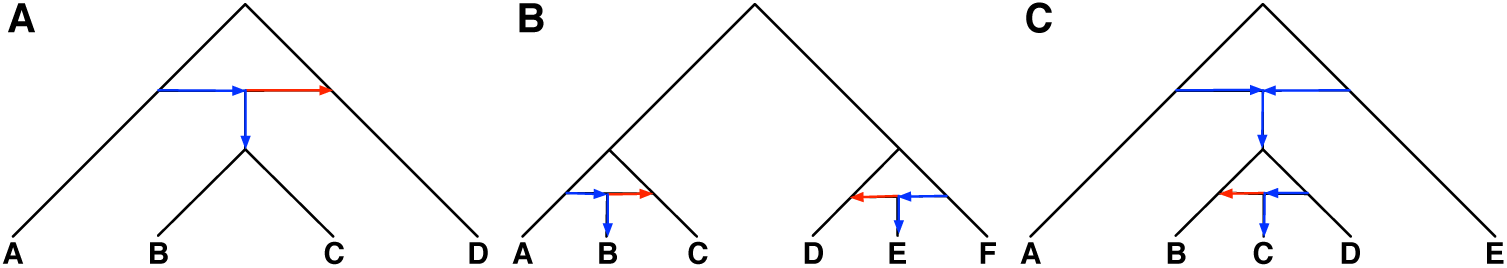
Networks inferred under the NCM model and did not match the true networks. (A) The network inferred from the data generated on the network of Scenario I and branch length setting 2. (B) The network inferred from the data generated on the network of Scenario II and both branch length settings. (C) The network inferred from the data generated on the network of Scenario III and branch length setting 2. Red arrows indicate the reticulations whose direction was inferred in the reverse order.

The only errors in the inferences were the direction of the reticulation edge (as highlighted with red arrows in Fig. 4). For Scenario I, while the true network has a reticulation from D to the ancestor of B and C, the inferred network has the reticulation in the opposite direction. For Scenario II, while the true network has a reticulation from C to B and from D to E, the inferred network has both reticulations in the opposite direction. For Scenario III, while the true network has a reticulation from B to C, the inferred network has the reticulation in the opposite direction. In other words, when inference under the NCM model was erroneous, it was always in terms of the direction of reticulations only. This error resulted in inflated error rates due to the sensitivity of the network topological distance measure used.

### 4.2 Analysis of Mosquito Data Set

We reanalyzed the *Anopheles* data of [3]. The data consist of one genome from each of the species *An*. *gambiae* (gam), *An*. *coluzzii* (col), *An*. *arabiensis* (ara), *An*. *quadriannulatus* (qua), *An*. *merus* (mer) and *An*. *melas* (mel). *An*. *christyi* serves as the outgroup for rooting the gene trees. We used the same set of gene trees from the autosomes that were used in the analyses of [19]. In particular, we used the same set of 669, 849, 564, and 709 loci from the 2L, 2R, 3L, and 3R chromosomes, respectively, and where 100 maximum likelihood bootstrap trees were inferred for each locus and used in the inference. These data are already available in DRYAD, entry doi:10.5061/dryad.tn47c. In [19], the phylogenetic network was inferred from the gene trees using the maximum likelihood method of [25]. The phylogenetic networks from the original study of [3], from the maximum likelihood analysis of [19], and the one we obtained under the NCM model are shown in Fig. 5.

**Figure 5:**
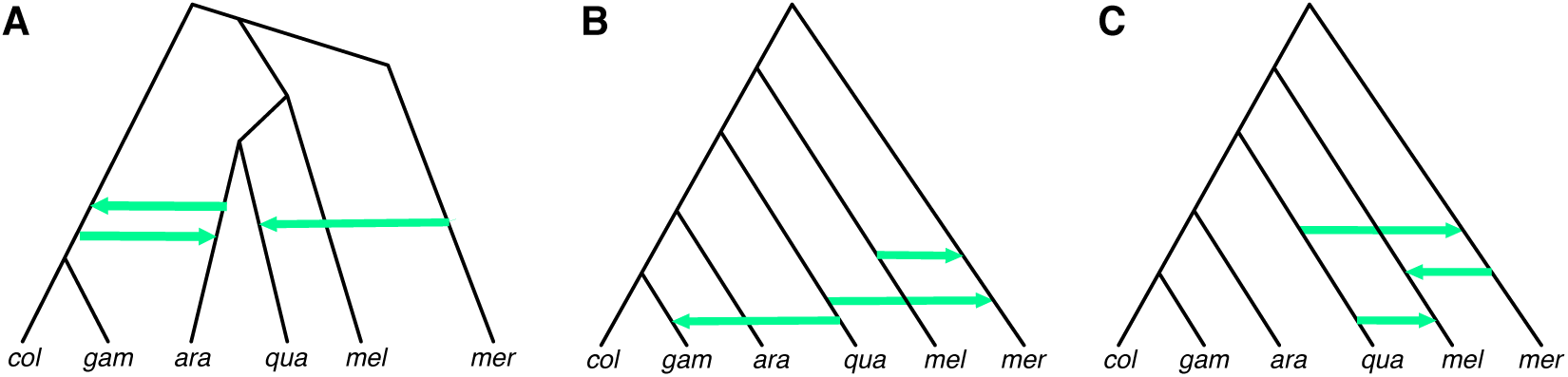
Networks for the mosquito data set. (A) The phylogenetic network reported in [3]. (B) The phylogenetic network analyzed using the maximum likelihood method of [25] and reported in [19]. (C) The phylogenetic network inferred under the NCM model.

As reported in [19], the likelihood of the network in Fig. 5(B) was much higher than that of the network reported by Fontaine *et al*. and shown in Fig. 5(A). The log-likelihood under the NCM model of the networks in Fig. 5(B) and 5(C) are −17520.49 and −16821.33, respectively. This demonstrates that the difference in the inferred networks is not due to limitations of the search procedure, but due to a better likelihood of the new inferred network over existing ones under the NCM model.

It is important to note both networks of Fig. 5(B) and 5(C) agree on the same underlying tree structure (the one obtained from the network by removing the arrows, and sometimes called the backbone tree). This tree disagrees with that reported by Fontaine et al. [3], and this disagreement was discussed in [19] (it has to do mainly with the fact that Fontaine *et al*. relied on information from the X chromosome to build a species tree and then augment it with postulated hybridization events). Our inferred network also agrees with that of [19] in terms of the *An*. *quadriannulatus* and *An*. *merus* hybridization. However, the *An*. *merus* and *An*. *melas* hybridization differ in terms of the direction of the reticulation (similar to the trend observed on the simulated data and discussed above), and the *An*. *quadriannulatus* and *An*. *melas* hybridization is not reported in [19].

There could be several reasons for the differences between the two networks of Fig. 5(B) and 5(C). The obvious one is that the two networks were inferred under two different models, one that a common mechanism of evolution underlies all loci and the other that assumes each locus has its own model. Second, as discussed in [22], (unpenalized) maximum likelihood cannot determine the correct number of reticulations. It could be that as more complex networks (ones with more than three reticulations) are inferred, the two analyses using the method of [25] and the one under the NCM model might converge onto the same network. These differences notwithstanding, inference under the NCM model recovered a very similar network, which makes it promising to explore the network space first without optimizing or sampling the continuous parameters, and then potentially follow up with a sampling phase to recover parameter values.

## 5 Discussion and Future Directions

In this paper, we introduced a no-common mechanism for phylogenomics, where the species phylogeny topology is the same across all loci, but the gene tree of every locus evolves under its own parameter (branch lengths and inheritance probabilities) settings. We implemented a maximum integrated likelihood function under this NCM model, assessed its accuracy and compared it to inferences under the parsimony MDC criterion on simulated data and the maximum likelihood inference on an empirical data set of mosquito genomes. We found that the inference produces very good results and when there is a disagreement, it is most often in terms of an incorrect direction (but not the placement) of reticulation edges. The main rationale behind developing such a mechanism is to allow for developing methods for efficiently exploring the species phylogeny space while focusing on traversing different topologies without the need for sampling or optimizing branch lengths and other continuous parameters.

Tuffley and Steel [18] showed that an NCM model is related to Fitch’s parsimony for character evolution in the following way. The maximum likelihood tree (or trees) under their NCM model is also a maximum parsimony tree under certain conditions. In light of this, we investigated whether a similar correspondence might hold for the NCM model for gene trees evolving down a species trees. Using the three gene trees of Fig. 6, and assuming equal frequencies of all three, we inferred the optimal tree under the MDC criterion as well as the optimal tree under the NCM model. Fig. 7 shows the results. As the results show, the two criteria in this case result in different optimal species trees. Furthermore, in this case, the optimal tree under the NCM model is not unique.

**Figure 6:**
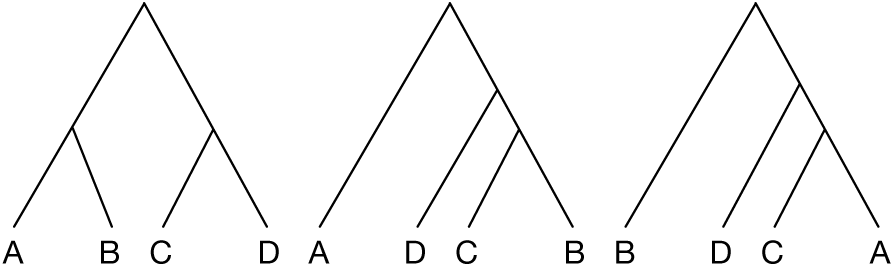
Three gene trees with equal frequencies that result in different optimal species trees under the MDC criterion and NCM model.

**Figure 7:**
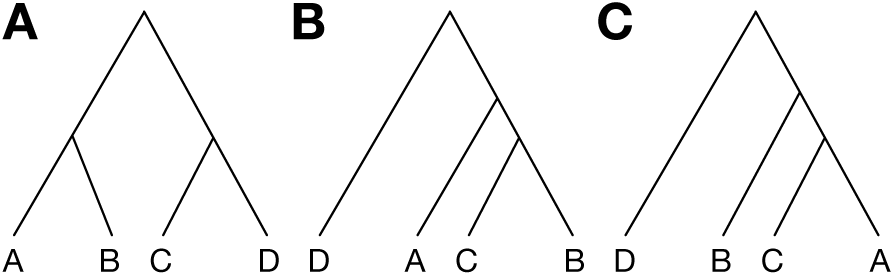
Optimal networks on the set of gene trees of Fig. 6. (A) Optimal species tree under the MDC criterion (has a cost of 4 extra lineages). The other two trees each have a cost of 5 extra lineages. (B) and Optimal species trees under the NCM model.

The work gives rise to several questions. First, under which conditions are optimal trees or networks identical under both the MDC criterion and the NCM model? Second, is there a parsimony criterion other than MDC under which the optimal species phylogeny is always identical to the optimal tree under the NCM model? Third, under what conditions, if any, is inference under the NCM model statistically consistent? Fourth, in light of the results above, an important question would be to explore why optimal networks under the NCM model often have reticulations in the opposite direction from those in optimal networks under the likelihood model with a common mechanism across all loci. Answering these and other questions will open up many research avenues in the area of phylogenomics.

Finally, while the simulations reported above were conducted on data generated under a common mechanism, it is important to conduct studies where different loci have different evolutionary parameters so as to mimic data coming from autosomes and sex chromosomes, as well as loci under selection or where duplication and loss could have played a role. In such cases, assuming a common mechanism underlying all loci is inappropriate and inference under an NCM model could provide more accurate results.

## Acknowledgments

The authors thank Jiafan Zhu for help with implementing the method and running the experiments. This work was supported in part by NSF grants DBI-1355998, CCF-1302179, CCF-1514177, CCF-1800723, and DMS-1547433.

